# Digital polarimetric second harmonic generation microscopy of partially oriented fiber structures

**DOI:** 10.1101/2022.08.23.504933

**Authors:** Mehdi Alizadeh, Serguei Krouglov, Virginijus Barzda

## Abstract

Second harmonic generation (SHG) in biological tissue originates predominantly from noncentrosymmetric fibrillar structures partially oriented within the focal volume (voxel) of a multiphoton excitation microscope. The study is aimed to elucidate fibrillar organization factors influencing SHG intensity, as well as achiral, R, and chiral, C, nonlinear susceptibility tensor component ratios. SHG response is calculated for various configurations of fibrils in a voxel using digital nonlinear microscope. The R and C ratios are calculated using linear incident and outgoing polarization states that simulate polarization-in polarization-out (PIPO) polarimetric measurements. The investigation shows strong SHG intensity dependence on parallel/antiparallel fiber organization. The R and C ratio is strongly influenced by the fiber chirality, tilting of the fibers out of image plane and crossing of the fibers. The study facilitates interpretation of polarimetric SHG microscopy images in terms of ultrastructural organization of fibers in the imaged structures.

**Statement of Significance:** Second harmonic generation microscopy is widely used for imaging non-centrosymmetric biological structures such as collagen. The ultrastructure of collagen can be determined with polarimetric SHG microscopy. The coherent nonlinear response of biological structures depends on the 3D orientations and positions of the collagen fibers in the focal volume of the microscope. Here, we show how different fiber organizations and 3D orientations in the focal volume can affect the polarimetric SHG responses. The results are important for understanding and interpreting images obtained with polarimetric SHG microscopy.

## 1 Introduction

Second harmonic generation (SHG) microscopy is an indispensable ultrastructure characterization technique for biological samples containing non-centrosymmetric fibrillar structures such as collagen (1–5), myosin (6, 7), or starch granules (8–10). The fibrillar structures are partially oriented in the biological samples, thus polarimetric SHG microscopy can be used to extract information about molecular organization in each focal volume of the imaged structures (3, 4, 6, 11–13). Various SHG polarimetry techniques have been used to reveal ultrastructure of biological samples (3, 8, 14–17). A complete Stokes-Mueller polarimetry in two dimensions (2D) can extract complex valued measurable nonlinear susceptibility tensor components (15, 16, 18). The reduced polarimetry techniques may employ linear incident polarization states (2, 11, 12, 14), or incident and outgoing linear polarizations (3, 19–23). Circular polarization states are also used for the in-image-plane orientation independent measurements, revealing the susceptibility ratios (8, 17).

The polarimetric nonlinear microscopy uses high numerical aperture (NA) objectives for fundamental beam focusing into a tiny voxel. The voxel volume is usually much larger than the diameter of a second harmonic active fibrillar structure, and many fibers with different arrangements can be present in the voxel (2). The hierarchical organization of fibrillar harmonophores in the biological samples can be divided into (i) short range fiber configurations, where phase differences between harmonophores can be neglected, and (ii) configurations separated by a fraction of wavelength of light where phase relations have to be considered. A numerical modeling has been previously presented to study interactions of laser light with nonlinear media in a focal volume of high NA objectives (24). This modeling showed that polarimetric SHG microscopy parameters, e.g. achiral susceptibility ratio, depends on the NA of excitation and collection objectives. In this work, the dependencies of achiral (R) and chiral (C) susceptibility tensor component ratios, as well as the effective in-image-plane orientation angle (*δ*) and out-of-imageplane tilt angle (*α*) of different fiber configurations appearing in a voxel are studied analytically and also by performing numerical modeling (24). The outcomes of simulations from this study can be used for interpreting the results of polarimetric SHG measurements of biological samples.

## 2 Materials and methods

The nonlinear response of fibrillar material is investigated using numerical modeling with digital nonlinear microscope (24). The phase relations of fundamental radiation and SHG from fibers located at different positions in the focal volume are taken into account. In this modeling, the angular spectrum representation is used for the focusing beam, where the incoming fundamental electric field that passes through the excitation objective changes its propagation direction corresponding to the lateral distance from the optical axis. The beam interacts with the material of a given geometrical configuration of fibers in the focal volume. The fibers have specified values of molecular nonlinear susceptibility tensor elements. A nonlinear polarization for arbitrary spatial distribution of fibrillar harmonophores within the voxel is generated and its harmonic radiation is collected with the collection microscope objective in the far field. The simulation results in an SHG intensity image of a back entrance aperture of the collection objective. SHG intensity is calculated by summing all pixel values in the image.

In this study, a linear polarization at different orientations is used for incoming fundamental radiation, and the linearly polarized SHG response is detected after passing the signal through a linear analyzer oriented at different angles in order to simulate polarization-in polarization-out (PIPO) SHG microscopy measurements. The digital PIPO microscope employs eight incident linear polarizations at orientations differing by 22.5°angles with respect to the Z laboratory axis. The resultant far field SHG electric field is calculated, and subjected to the polarization state analyzer for a set of eight orientations of linear polarizer at different angles rotated by steps of 22.5°from the Z laboratory coordinate axis. The obtained SHG intensities with the set of incident and outgoing polarization states are used for modeling with the nonlinear digital microscope (24). The modeling outcome provides with polarization independent SHG intensity values (average over all linear polarization states), as well as achiral, R, and chiral, C, nonlinear susceptibility ratio values of the fibrillar structures within the focal volume. The NA of both excitation and collection objectives is set to 0.8 in all cases. The fiber diameters are all set to 100 nm, while the voxel itself has 1*µm* lateral and 2.4*µm* axial full width at half maximum (FWHM) fundamental intensity profile. The wavelength of the fundamental radiation is set to 1028 nm. It is assumed that R ratio is real and C ratio is complex in all modeling cases (15). The molecular susceptibility tensor elements 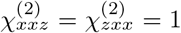 are set for all samples. The tensor elements of 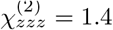 and 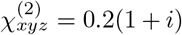 represent collagen and 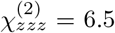 and 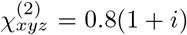 are chosen for TPPS_4_ fibrillar aggregates.

The SHG intensity obtained with PIPO measurements for each pixel containing *C*_6_ symmetry fibers is as follows (3):

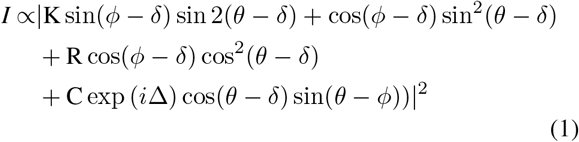

where 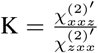 and K=1 is assumed for off resonance excitation conditions, 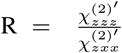 is the achiral susceptibility ratio, 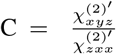 is the chiral susceptibility ratio, *δ* is the in-image-plane effective orientation angle of the fibers, *θ* is the incoming linear polarization orientation angle, and *ϕ* is the outgoing linear polarization orientation angle. The chiral susceptibility is assumed to be complex valued with a phase retardancy Δ between achiral and chiral susceptibility components (15, 25, 26). A schematic picture of a fiber in the focal volume is presented in Fig. 1. The sample plane is located in the XZ plane of the laboratory coordinate system and Y is the beam propagation direction. The molecular coordinate system of the fiber has z coordinate along the fiber axis, and x, y perpendicular to the fiber. The prime of the susceptibility elements indicates the implicit dependence on the out of plane tilt angle *α* and can be expressed in terms of the susceptibility tensor elements in the molecular frame as follows (3):

**Figure 1:**
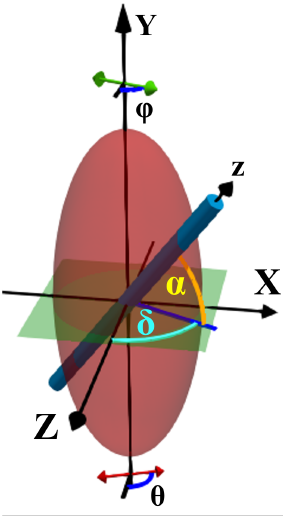
Laboratory coordinate system for a cylindrical fiber with *C*_6_ symmetry in the voxel. The fiber has a projection in the image plane (XZ) with an in-plane *δ* orientation angle from Z axis and a tilt angle of *α* out of the image plane. *θ* denotes the incoming and *ϕ* denotes the outgoing polarization orientation angle measured with respect to the laboratory Z axis. The molecular z axis is along the fiber axis.

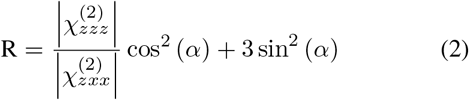

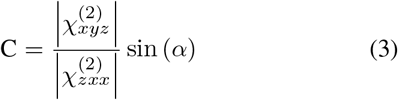

## 3 Results

Modeling is conducted for chiral fibers arbitrarily organized in a focal volume of high NA microscope objective. The fibers can be divided into two categories with 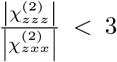 and 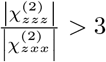, based on their molecular achiral susceptibility ratio. The ratio 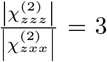 has a critical value that gives no tilt angle *α* dependence for K=1 (see eq (2)) (17, 27). For modeling of collagen fibers, a molecular susceptibility ratio of 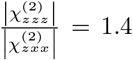 is assumed. For an example of fibers with achiral molecular susceptibility ratio above 3, fibrillar TPPS_4_ aggregates with 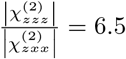 are considered (28). Main fiber configurations affecting the SHG intensity, *I*, R and C ratios in the focal volume are presented in the following sections.

### 3.1 A single fiber

#### 3.1.1 A single fiber in the image plane

The polarimetric SHG response is modeled for one fiber oriented parallel to the image plane (*α* = 0). When displacing the fiber axially or laterally by the distance *l* from the center, the total SHG intensity follows the point spread function (PSF) profile, which is 2.4 times broader in axial than lateral directions (Fig. 2 b and c). In all plots, SHG intensities are normalized to the maximum intensity of one fiber located in the center of the focal volume and oriented parallel to the image plane.

**Figure 2:**
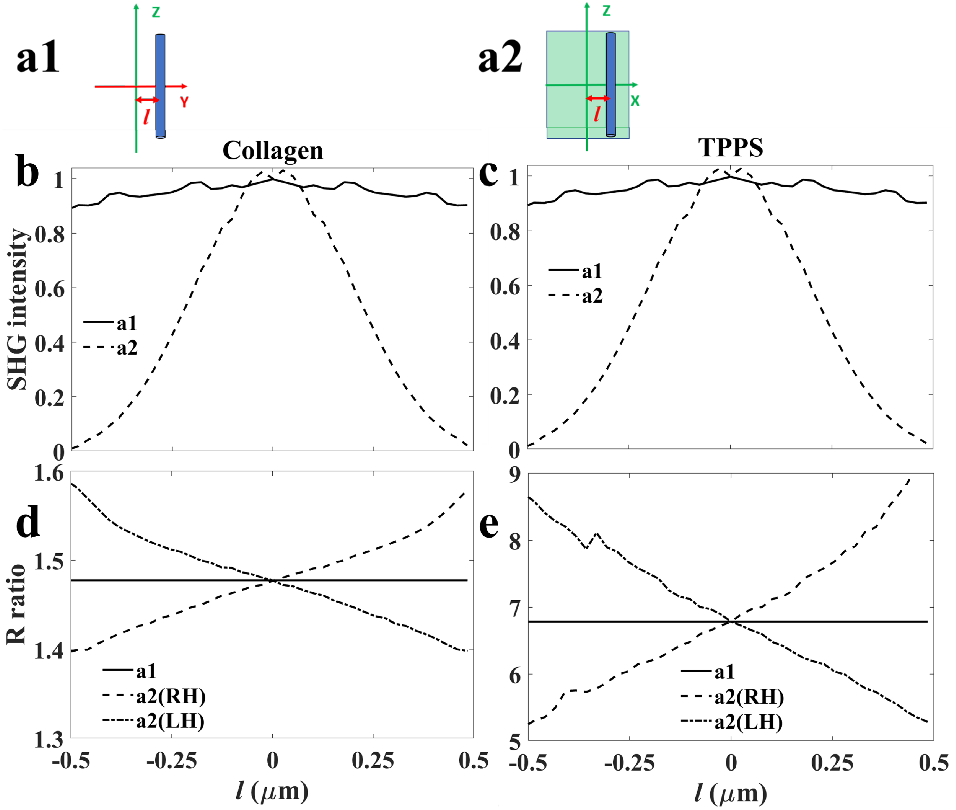
Normalized SHG intensity, *I* (b, c), and R ratio (d, e) dependency on the axial and lateral displacement, *l*, of a fiber from the center of focal volume for collagen (b, d) and TPPS_4_ aggregates (c, e). The configuration of a fiber in the focal volume is shown for axial (a1) and lateral (a2) displacements. The R ratio dependency is shown for axial and lateral displacements of right-handed (RH) and left-handed (LH) fibers (d, e), as indicated in the figure legends.

The polarimetric response of the fibers under focusing beam conditions results in increase of the observed achiral susceptibility ratio (24) to R=1.47 for collagen and R=6.77 for TPPS_4_ fibers. The modeled R ratio value does not change with the displacement in the focal volume when real-valued molecular susceptibility tensor elements are assumed. Interestingly, when complex valued chiral susceptibility 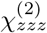 is assumed, the left side lateral displacement of a right-handed fiber leads to R ratio decrease, while the right side displacement gives higher R ratio. The effect is opposite for left-handed structure (dashed and dash-dotted lines in Fig. 2 d and e). The structure is considered to be right-handed when 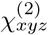 and 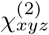 have the same sign; for opposite signs the structure is left-handed. The change in R with lateral displacement is observed due to strongly focused fundamental beams, where circular polarization component of electric field appears in the focal volume and evokes the imaginary part of 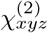. In contrast, R ratio remains unchanged when the fiber is moved axially (see the solid lines in Fig. 2 d and e). C ratio remains C=0 for all displacements from the center for fiber orientation parallel to the image plane (eq (3)).

#### 3.1.2 A single fiber tilted out of the image plane

Tilting fibers out of the image plane is one of the major effects influencing SHG intensity, *I*, R and C ratios. The SHG intensity increases with the tilt for collagen and decreases for TPPS_4_ aggregates (Fig. 3 b and c, see eq (1)-(3)). The SHG intensity from the tilted fiber is normalized to the intensity of a single fiber located in the center of the voxel and parallel to the image plane. The R ratio increases with the tilt for collagen and decreases for TPPS_4_ (Fig. 3 d and e), as previously reported (see eq (2)) (19).

**Figure 3:**
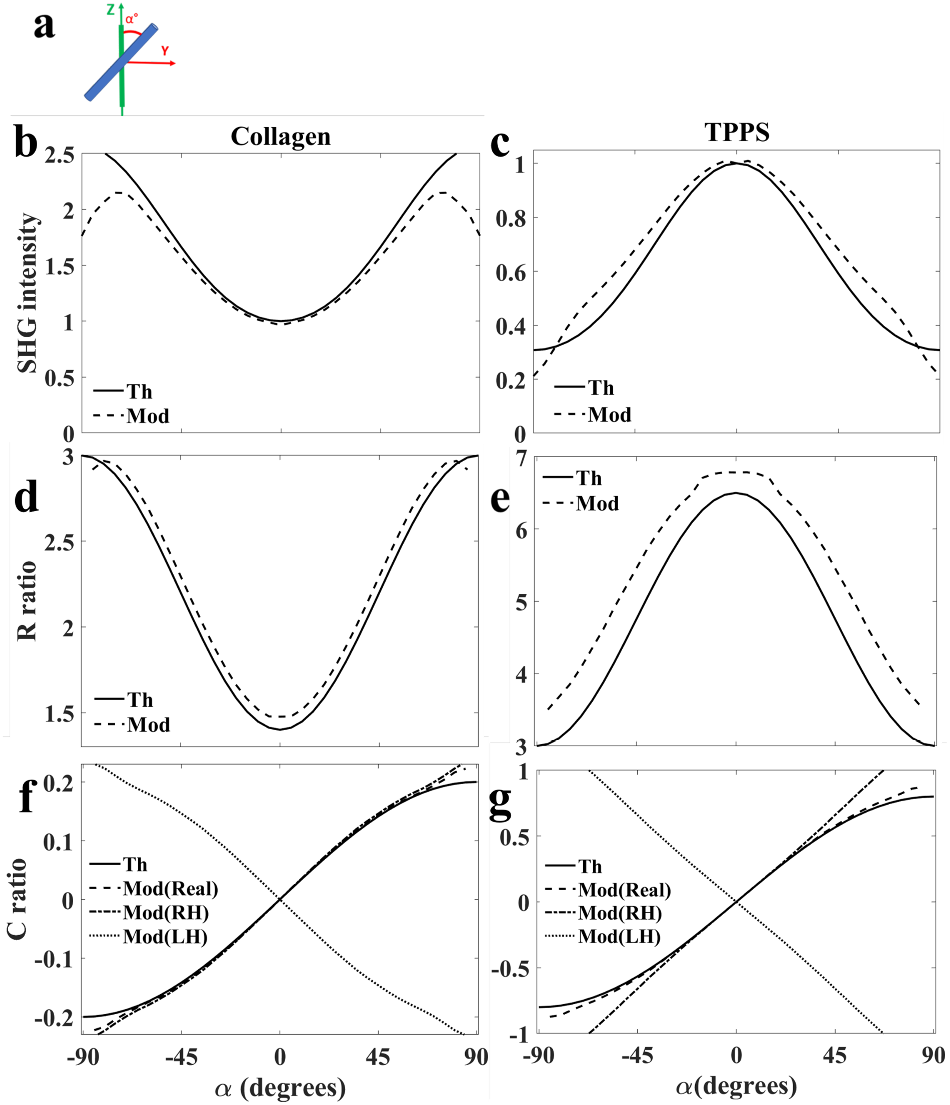
The normalized SHG intensity, *I* (b, c), R (d, e) and C (f, g) ratio dependence on the tilt angle *α* out of image plane for collagen (b, d, f) and TPPS_4_aggregates (c, e, g). The fiber configuration is shown in (a). The theory curves (Th, solid line) and modeling (Mod, dash/dotted line) were obtained with R=1.4 for collagen (d) and R=6.5 for TPPS_4_ aggregates (e). C ratio variations (f, g) are calculated from eq (3) (solid line) and numerically modeled with real valued 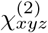 (dashed line), and complex valued 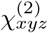 for righthanded (RH, dash-dotted line) and left-handed (LH, dotted line) fibers.

The numerical modeling of C ratio dependence on the tilt angle follows well eq (3) for real-valued 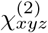 (Fig. 3 f and g). The sign of C ratio changes due to clockwise or counterclockwise tilt of the fiber around X-axis (eq (3)). When the complex values of 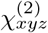 are assumed, some deviations from sinusoidal function appear for both, collagen and TPPS_4_ aggregates (Fig. 3 f and g). The presented deviation from eq (3) is higher for TPPS_4_ aggregates than collagen, because C_Collagen_ = 0.2(1+*i*) is assumed, while higher value of C_TPPS_ = 0.8(1 + *i*) is taken in our modeling. Fig. 3 f and g shows that switching the handedness of the fiber changes the sign of C ratio, while R ratio sign is unchanged.

### 3.2 Parallel and anti-parallel fibers

#### 3.2.1 Two parallel and anti-parallel fibers

##### 3.2.1.1 Two fibers in the image plane

Configuration of parallel fibers oriented along the image plane results in high SHG signal intensity, especially for fibers stacked axially in the focal volume. Fig. 4 b and c show SHG intensity dependence on lateral displacement 2*l* of fibers from each other, symmetrically from the center of the focal volume. The intensity values are normalized to a single fiber centrally located in the focal volume. When two adjacent antiparallel fibers are present in the focal volume the SHG intensity is significantly reduced (Fig. 4 b and c). The resultant SHG intensities show a coherent summation of the signals from the fibers. Less than 3% remains of the intensity of a single fiber in the center for axially adjacent antiparallel fibers, and the SHG intensity gradually increases with the fiber separation. For the laterally adjacent fibers, the intensity is slightly higher and increases to a maximum value that is slightly higher than for one fiber, and then the intensity decreases again with further separation between the two fibers. This phenomena can be used for nanopositioning experiments between the two SHG emitters in the focal volume. The intensity dependence with the displacement is very similar for both, collagen and TPPS_4_. The achiral susceptibility R ratio for the parallel fibers is similar to an individual fiber in the focal volume, although there are small modifications for some configurations (see Fig. 4 d and e). The trends of R with separation are very similar for parallel collagen and TPPS_4_ fibers. When two antiparallel fibers are symmetrically laterally displaced from each other the R ratio is low at small separations and increases when fibers get further apart (Fig. 4 d and e). The trends for lateral separation again are similar for collagen and TPPS_4_ aggregates. The R values cannot be accurately modelled for axial separation at very close proximity due to weak SHG signal. Further separation of two antiparallel collagen fibers axially results in low R ratio value, which gradually increases as shown by the dashed line in Fig. 4 d. When two antiparallel TPPS_4_ fibers are displaced axially R ratio starts with high values and then decreases eventually plateauing at R=6.9 (Fig. 4 e). Note, that C=0 for the fibers parallel to the image plane.

**Figure 4:**
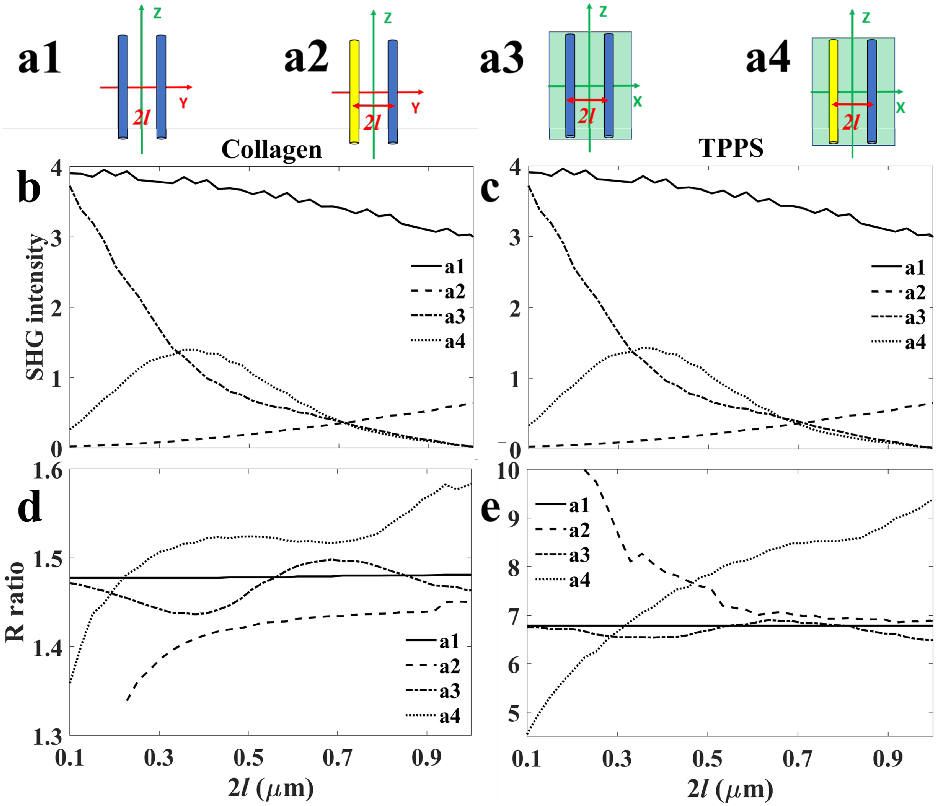
SHG intensity (b, c) and R-ratio (d, e) dependence on the separation distance 2*l* between the collagen (b, d) and TPPS_4_ fibers (c, e), for parallel and antiparallel fiber configurations located parallel to the image plane. The fibers are shown in YZ image plane (a1, a2), or in XZ plane (a3, a4) with Y coordinate along the light propagation direction. Fibers in blue and yellow designate opposite polarity. The curve styles and corresponding fiber configurations (a1-a4) are indicated in the figure legends.

##### 3.2.1.2 Two fibers tilted out of the image plane

The results of tilting two parallel and antiparallel fibers out of the image plane are presented in Fig. 5 (see panel a1a4 for respective configurations). Fig. 5 b and c show that the normalized total SHG intensities increase for collagen and decrease for TPPS_4_ by tilting axially (solid line) and laterally (dash-dotted line) arranged parallel fibers. The SHG intensity diminishes for antiparallel fibers, and, when tilted, both collagen and TPPS_4_ fibers show some intensity increase as presented by dashed and dotted curves.

**Figure 5:**
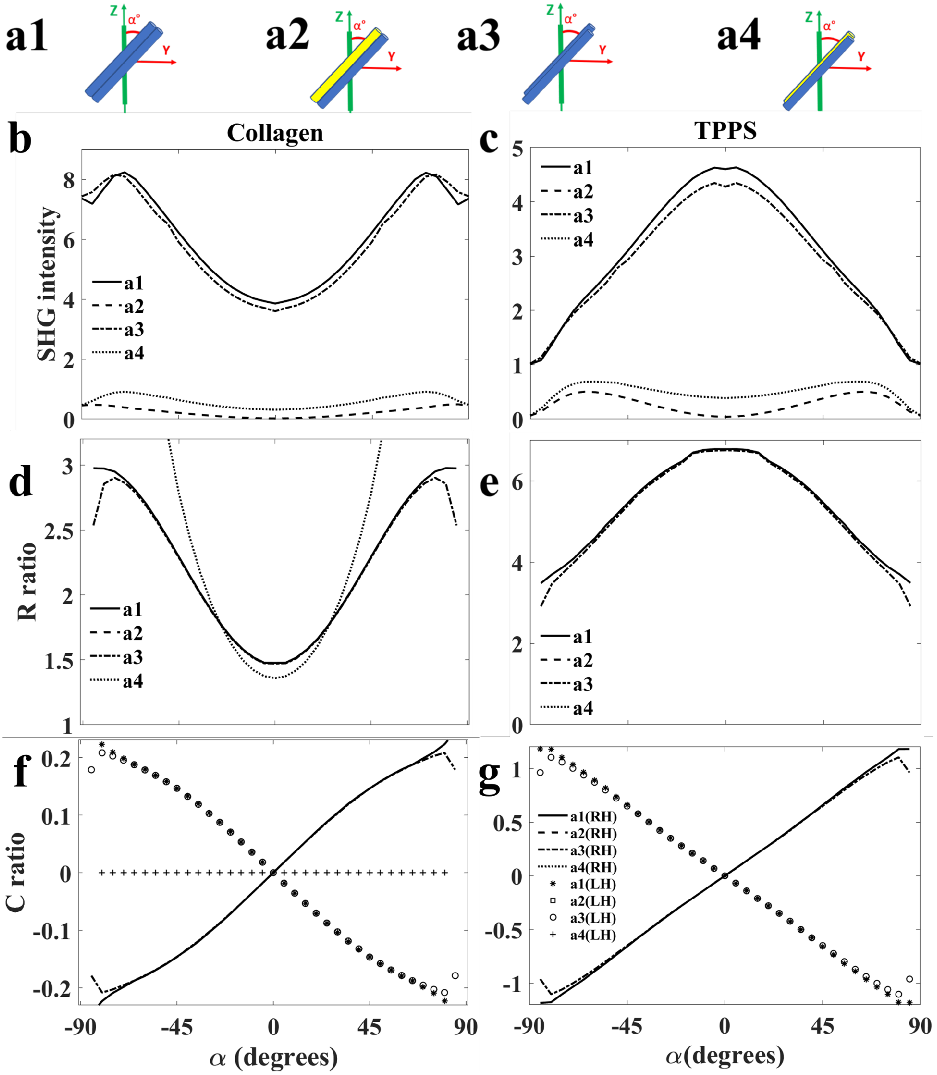
SHG intensity (b, c), R (d, e) and C (f, g) ratio dependence on tilt angle *α* for axially and laterally parallel (a1, a3 blue) and antiparallel (a2, a4 blue-yellow) fibers, respectively, for collagen (b, d, f) and TPPS_4_ aggregates (c, e, g). The curve styles of axially and laterally parallel and antiparallel configurations are indicated in the figure legends for left-handed (LH) and right-handed (RH) fibers.

The R ratio increases with tilt angle *α* for parallel collagen and decreases for TPPS_4_ fibers (Fig. 5 d and e). For antiparallel configuration, only laterally stacked collagen fibers could be modeled and gave lower initial R ratio and steeper dependence with the tilt (Fig. 5 d).

The C ratio of parallel fibers followed similar trend with tilt angle as for a single fiber (Fig 5, f and g). The C ratio with tilt could not be calculated for the antiparallel fibers for both, collagen and TPPS_4_. Furthermore, changing the handedness of fibers doesn’t affect the results of *I* and R, but results in changing the sign of C ratio, as shown by asterisk and circle lines in panels f and g in Fig. 5.

#### 3.2.2 A bundle of parallel and anti-parallel fibers

##### 3.2.2.1 A bundle in the image plane

Several fibers can assemble into a bundle resembling biological structures (2, 29–31). A fiber bundle encompasses axial and lateral configuration dependent nonlinear effects presented above. Fig. 6, a1-a8 presents fiber bundles with five fibers arranged one in the center, two in the left and right positions and two in the front and back positions of the central fiber. The fibers oriented in one direction are colored in blue and flipped fibers are colored in yellow. For each configuration, the normalized SHG intensities at *α* = 0 are very similar when comparing collagen and TPPS_4_ bundles (Fig. 6, b and c). The variation in SHG intensity is determined by coherent summation of SHG signals from individual fibers. The minimum intensity is achieved at the half of flipped fibers. This effect appears in stripy tendon structure (30, 31), where high and low SHG intensity stripes have mostly parallel and antiparallel fibers, respectively. The R values have small variation between the configurations at *α* = 0 (Fig 6, d and e). The R ratio appears around R=1.47 for collagen and R=6.75 for TPPS_4_ fiber bundle. The antiparallel structures may slightly modify the R ratio that increase a spread of R values in the experiments (32, 33). The C ratio is 0 for the fiber bundles with *α*=0.

**Figure 6:**
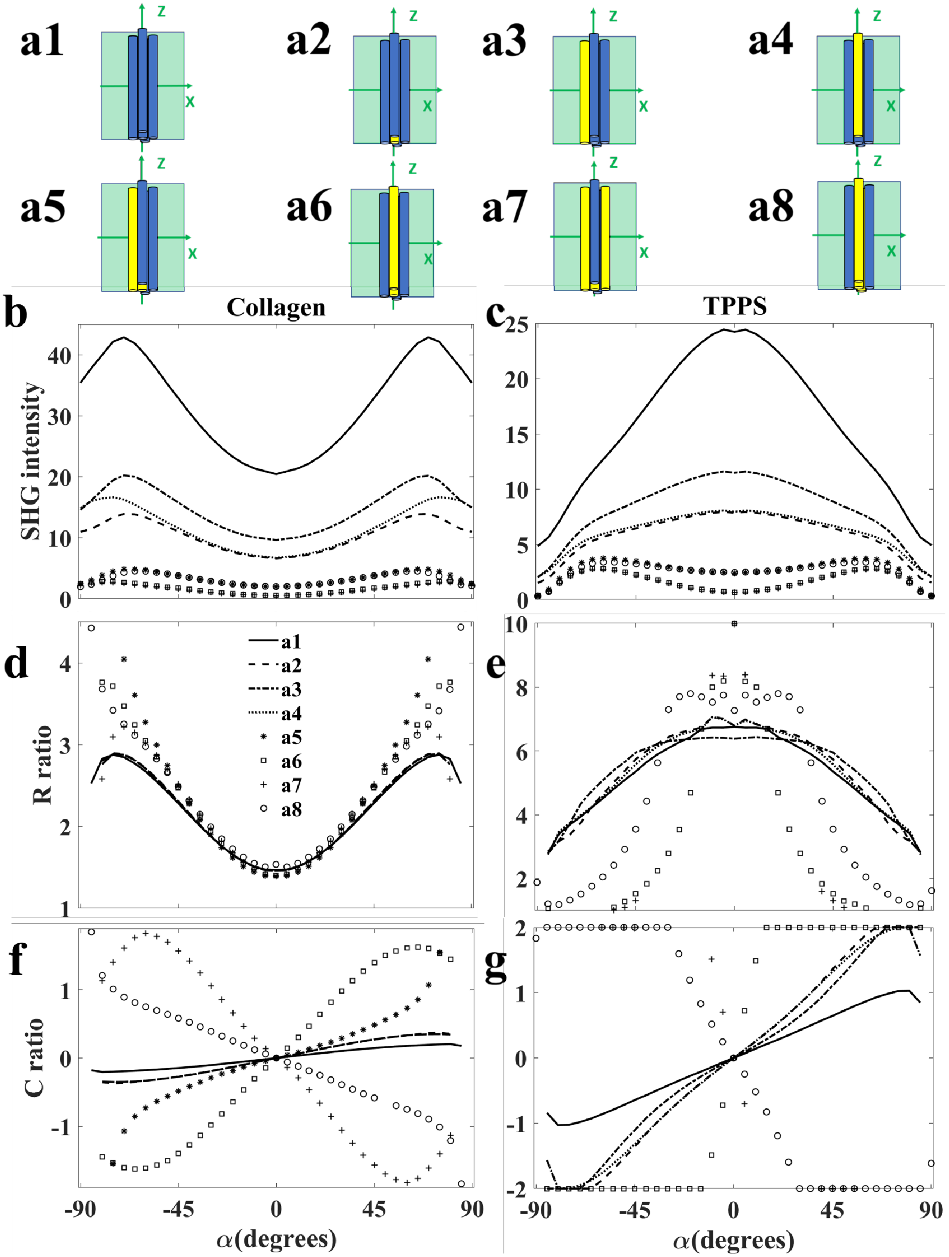
SHG intensity (b, c), R ratio (d, e) and C ratio (f, g) dependence on bundle structure and tilt angle *α* for axially parallel (blue, (a1)) and antiparallel (blue, and flipped fibers in yellow, (a2-a8)) fibers of collagen (b, d, f) and TPPS_4_ aggregates (c, e, g). The bundle configuration is as follows: five parallel fibers (a1), the central fiber is flipped (a2), the fiber on the left is flipped (a3), the fiber on the top is flipped (a4), the central fiber and the fiber on the left are flipped (a5), the central fiber and the fiber on the top are flipped (a6), three lateral fibers are flipped (a7), three axial fibers (along the axial direction of beam propagation) are flipped (a8). The curves corresponding to a1-a8 configurations are indicated in the figure legend on (d).

##### 3.2.2.2 Tilting a bundle out of image plane

The bundles of fibers are often found tilted out of the image plane. The results of the PIPO digital microscope for tilting bundles of five parallel and antiparallel fiber configurations (Fig. 6, a1-a8) are presented in Fig. 6, b, d and f for collagen and Fig 6 c, e and g for TPPS_4_ aggregates.

When tilting the bundles out of image plane, the SHG intensity increases for collagen and decreases for TPPS_4_ aggregates. The exception is for TPPS_4_ aggregates with close to balance of parallel and antiparallel fibers when two or three out of five fibers are flipped (cases a5-a8), resulting in low SHG intensity.

The curves in Fig. 6, d and e show the R ratio change with tilt for different bundle structures, which resembles behaviour of a single fiber. The R ratio increases when a bundle of collagen fibers is tilted, and this ratio increases faster when more than one fiber is flipped in the bundle. The TPPS_4_ fibrillar bundles show opposite trend with decrease in R values for increasing tilts out of the image plane.

Fig. 6, f and g show the behavior of C ratio for the bundles of fibers tilted out of the image plane. C behavior for antiparallel structures have a high deviation from sinusoidal curve when more fibers are flipped. The deviation from sinusoidal behavior increases until 50% of fibers are parallel and 50% are flipped. At higher than 50% flip, the C ratio dependence changes the sign, and when all the fibers are flipped C ratio behavior becomes sinusoidal, but with the negative sign. The 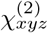 tensor elements are complex for all configurations in Fig. 6. Note, when bundles contain about 50% fibers flipped, the SHG intensity is low, and R and C ratios deviate significantly from eq (2) and (3), respectively.

### 3.3 Crossed fibers in 2D oriented structures

#### 3.3.1 Two crossed fibers

##### 3.3.1.1 Two crossed fibers in the image plane

A configuration of crossing fibers significantly influences the results of polarimetric nonlinear microscopy (2). Fig. 7 shows the normalized SHG intensity, R ratio and C ratio variations with respect to the crossing angle *σ* between two fibers situated within the image plane, as it is shown schematically in Fig. 7, a. In both samples, the SHG intensity decreases by increasing crossing angle *σ* between the two fibers (Fig. 7, b and c). The modeling shows that for the structures with *R <* 3, the R ratio increases with increasing *σ* (Fig. 7, d). This effect is opposite for *R >* 3 (Fig. 7, e). The C ratio is close to 0 when the crossing fibers are in the image plane (Fig. 7, f and g). Changing the handedness of individual fibers does not change PIPO parameters and, therefore, respective curves of *I*, R, and C overlap (solid and dashed curves, Fig. 7). There is a slight difference in C ratio at large *σ* for clockwise (right-handed) and counter clockwise (left-handed) crossing fiber arrangements (see Fig. 7, f).

**Figure 7:**
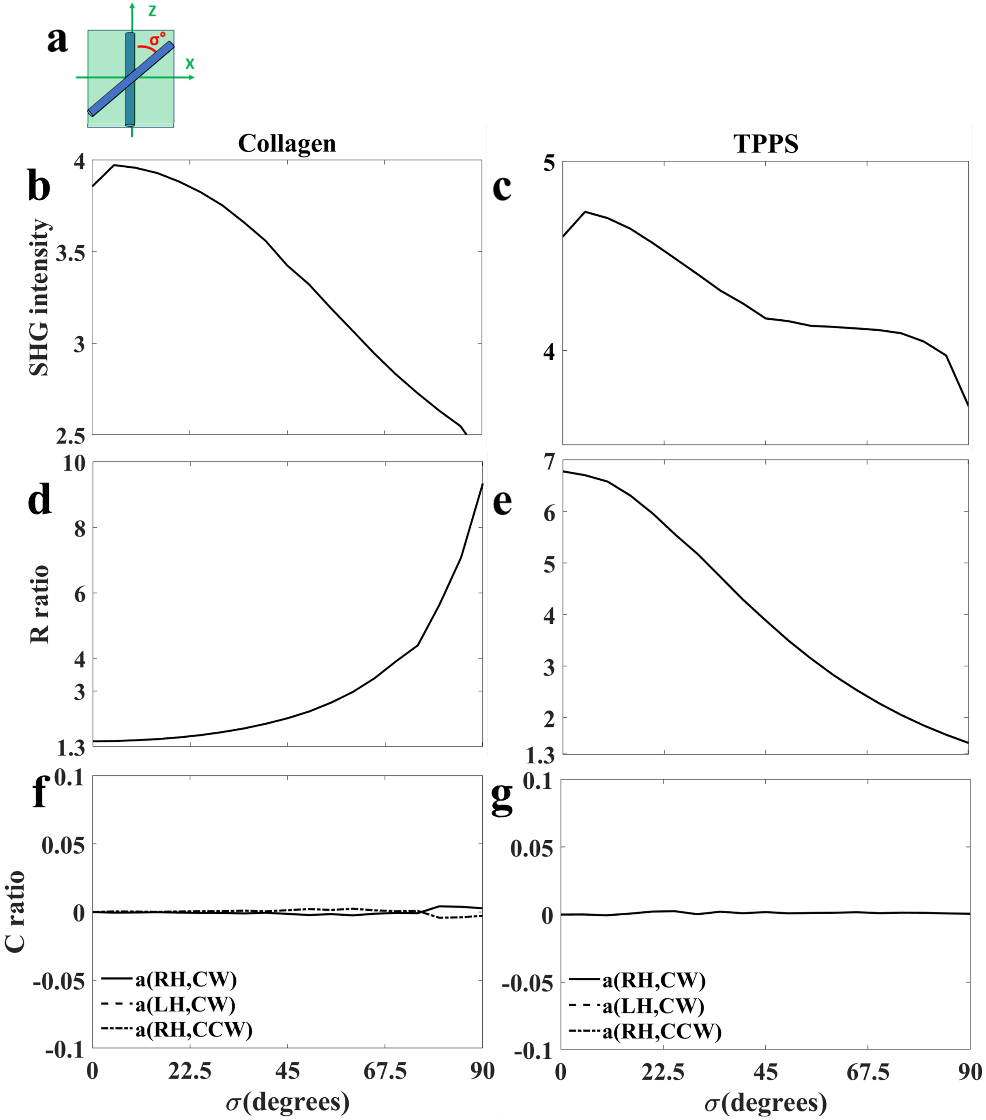
Numerical modeling of two crossing fibers at angle *σ*, oriented within the image plane (schematic picture in (a)). Normalized SHG intensity (b and c), R ratio (d and e), and C ratio (f and g) dependence on crossing angle *σ* for collagen (b, d, f) and TPPS_4_ fibers (c, e, g). Right-hand (RH), or lefthand (LH) fibers constitute the clockwise (CW) or counter clockwise (CCW) crossing shown in the figure legend.

##### 3.3.1.2 Two crossed fibers tilted out of the image plane

The tilt angle effects on SHG intensity, R, and C ratios are examined for crossed fiber structures at *σ*=20°and *σ*=45°, that lead to different initial values of *I* and R at *α*=0 (see Fig, 8). SHG intensity increases for collagen and decreases for TPPS_4_ with the tilt angle *α* (Fig. 8, b and c). The R ratio increases with *α* for collagen and decreases for TPPS_4_ fibers (Fig. 8, d and e). The C ratio has typical sin (*α*) dependence on tilt angle (Fig. 8, f and g). The C ratio changes sign for opposite chirality fibers.

**Figure 8:**
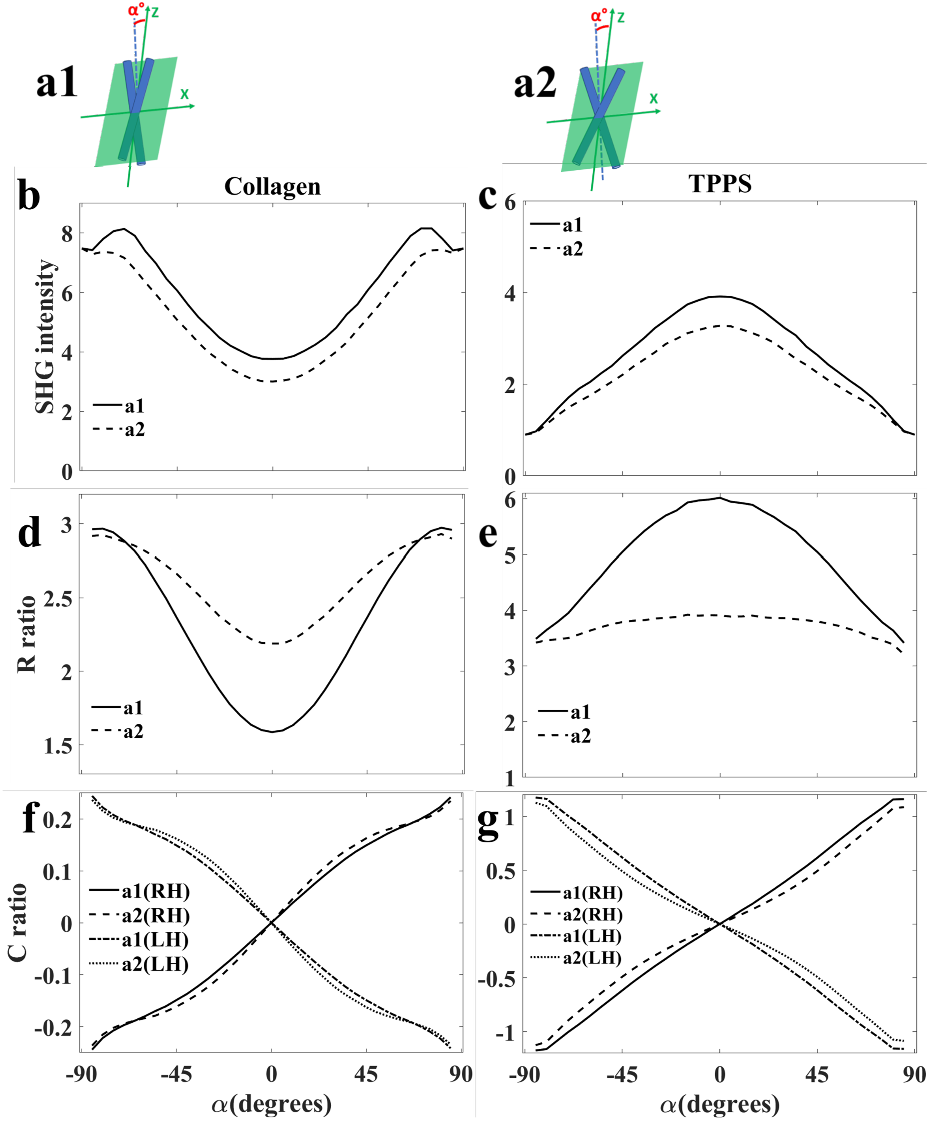
Numerical modeling of two crossing fibers tilted out of the image plane. Schematic picture of two fibers crossed at *σ*=20°(a1) and *σ*= 45°(a2) and tilted out of the image plane. Normalized SHG intensity (b and c), R ratio (d and e), and C ratio (f and g) variations with tilt for collagen (b, d, f) and TPPS_4_ aggregates (c, e, g). The sign of C ratio changes by switching individual fibers from right-handed (RH) to lefthanded (LH) structure. The fiber configuration information for each curve is indicated in the figure legends.

##### 3.3.1.3 Two crossed fibers in the plane perpendicular to the image plane

Fibers can cross when oriented in the plane along beam propagation direction. Fig. 9, a1 and a2 show two configurations of crossing fibers in the axially oriented plane; one is with fibers symmetrically oppositely tilted out of the image plane (Fig. 9, a1), and the other configuration shows the tilting of one fiber while the other remains in the image plane, resulting in tilt of effective axis of crossed fibers by 0.5*α* (Fig. 9, a2). The SHG intensity increases for collagen and decreases for TPPS_4_ fibers (Fig 9, b, c, respectively). The R ratio increases with increasing fibers tilt angle for collagen and decreases for TPPS_4_ fibers (Fig. 9, d and e, respectively). The C ratio remains close to 0 for the symmetric tilt of the fibers (see solid curves in Fig. 9, f and g) since the symmetrical opposite tilts in the focal volume cancel each other’s effect. The tilting of one fiber while other keeping parallel to the image plane results in the same behaviour of C ratio as tilting of one effective fiber by 0.5*α*.

**Figure 9:**
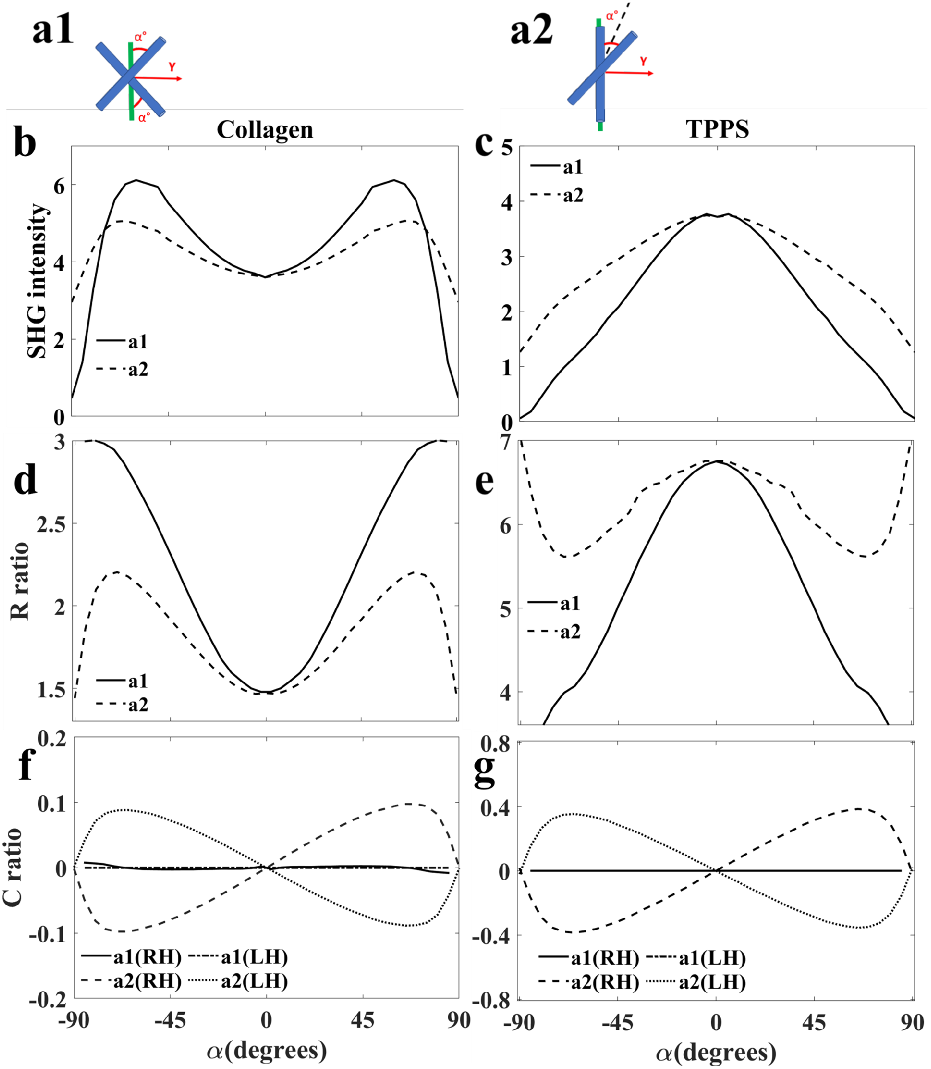
Numerical modeling of axially crossing fibers: Schematic picture of two crossed fibers symmetrically oppositely tilted out of the image plane (a1), and tilted one fiber out of image plane while the other remains in the image plane (a2). Normalized SHG intensity (b and c), R ratio (d and e), and C ratio (f and g) variations for collagen (b, d, f) and TPPS_4_ aggregates (c, e, g). The sign of C ratio changes by switching the individual fibers from right-handed (RH) to lefthanded (LH) structure. The fiber configuration information for each curve is indicated in the figure legends.

#### 3.3.2 Crossing flat structures, plywood geometry

##### 3.3.2.1 Crossed flat structures in the image plane

Parallel fiber layers may form crossed lamellar structures of plywood geometry appearing, for example, in trabecular lamellar bone (34). Fig. 10 shows PIPO digital microscopy results of crossing plywood structures. The plywood structure is composed of two layers of 3 flat fibers crossing at an effective angle *σ* (Fig. 10, a1). When the flat structures are parallel to the image plane (*α* = 0) the total SHG intensity decreases for both, collagen and TPPS_4_ aggregates with increasing *σ*. The R ratio increases for collagen and decreases for TPPS_4_ with increasing *σ* (Fig. 10, d and e). Fig. 10, a2 shows an alternating antiparallel plywood structure and Fig. 10, a3 shows a structure with opposite polarity configuration. The SHG intensity *I* decreases dramatically for antiparallel fibers and both R, and C ratios have the same behaviours as for plywood with parallel fibers. Changing the handedness of individual fibers doesn’t change the *I* and R results, and C ratio is also not affected due to in-plane orientation of the fibers, as shown by dotted, asterisk, and square mark lines in Fig. 10, f and g.

**Figure 10:**
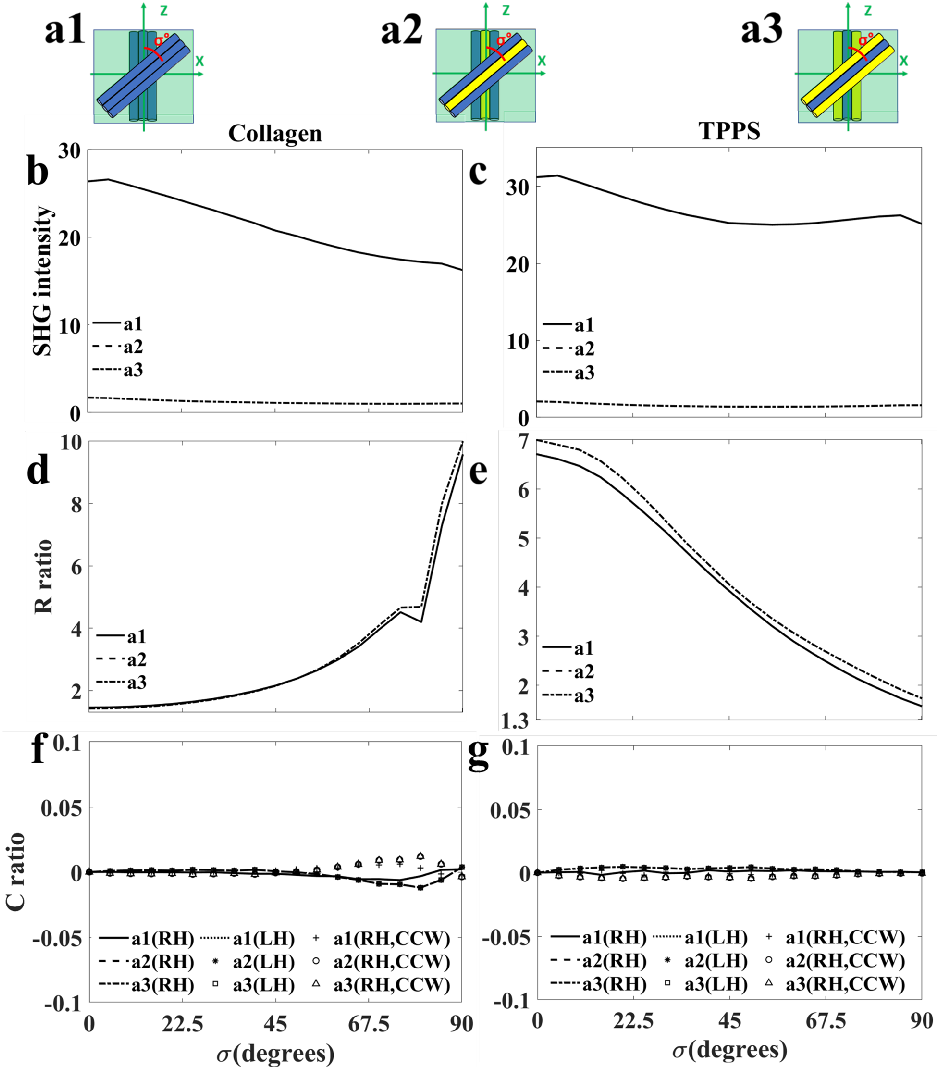
Numerical modeling of laterally crossed plywood structures at different crossing angles, *σ*. Schematic picture of crossing layers of parallel fibers (a1), alternating antiparallel fiber layers (a2) and opposite alternating antiparallel fiber layers crossed in the plywood structure (a3). SHG intensity *I* (b, c), R ratio (d, e) and C ratio (f, g) changes with *σ* for collagen (b, d, f) and TPPS_4_ aggregates (c, e, g).

The plywood geometry can result in right or left-handed macrochiral structure if the second fiber layer is turned clockwise or counterclockwise with respect to the orientation of fibers in the first layer. Changing the handedness of the crossing structure (macrochirality) leads to the same values for *I* and R, but C ratio changes the sign at large crossing angles, higher than 50°(Fig. 10, f). By increasing axial distance between the two layers the SHG intensity decreases slightly, but the R and C variation with *σ* remains the same.

##### 3.3.2.2 Tilting the plane of crossed layers

Tilting a plywood structure out of the image plane (Fig. 11) has an effect similar to the case of tilting two crossed fibers (Fig. 8). The configuration of crossed layers with parallel fibers is shown in Fig. 11, a1, and the structures with alternating polarity layers are presented in panels a2 and a3. The SHG intensity increases with tilt for collagen and decreases for TPPS_4_ aggregates, when parallel fibers in each layer are considered (a1 configuration). For alternating polarity layers, the *I* is dramatically decreased, and with tilt it is first increased, and at larger tilts decreased for both collagen and TPPS_4_ (dashed and dashed-dotted curves in Fig. 11 b and c). The R ratio for collagen increases with tilt for parallel layer configuration (Fig. 11, a1), and when the central fibers are flipped in both layers (Fig. 11, a2 and a3) the R ratio increases faster than for non-flipped collagen structure and reaches the limit of R=3 at a smaller *α* tilt angle. The R ratio for TPPS_4_ fibers decreases with tilt in a1 configuration, but first increases and then decreases with tilt for a2 and a3 configurations.

**Figure 11:**
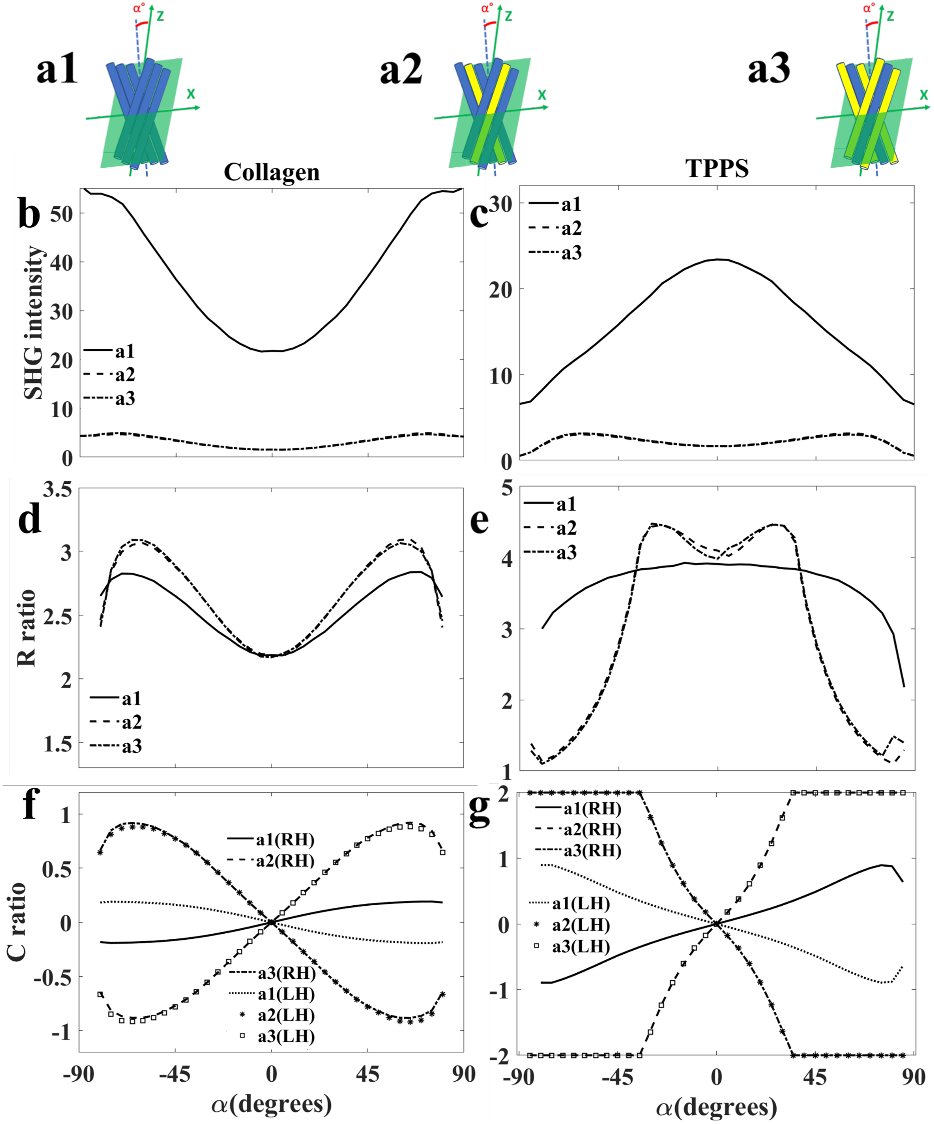
SHG polarimetric response from tilted plywood structures of crossing fiber layers at 45°and tilted by angle *α*. The fibers are arranged parallel (a1) or alternative antiparallel (a2, a3) in the layers. Normalized SHG intensity (b, c), R ratio (d and e), and C ratio (f, g) changes for collagen (b, d, f) and TPPS_4_ aggregates (c, e, g). The figure legends indicate structural configurations in (a), for the left handed (LH) and right handed (RH) constituting fibers (f, g).

The C ratio has typical sinusoidal behaviour for parallel fibers (a1) (Fig. 11, f and g). For antiparallel configurations (a2, a3), the C ratio with tilt is no longer sinusoidal and increases faster to the maximum for both antiparallel plywood arrangements. If the alternating arrangement is changed from a2 to a3 configuration (Fig. 11) the sign of the C ratio dependence on *α* flips for both, collagen and TPPS_4_ fibers. Changing the handedness of the fibers results in the sign flip of the C ratio curves (Fig. 11 f and g). The C ratio of TPPS_4_ fibers behaves very similar with the tilt as for the case of collagen, but the effect of deviation from sinusoidal dependence is much stronger in TPPS_4_ fibers (compare panels f and g in Fig. 11).

##### 3.3.2.3 Axially crossed plywood structures in the plane perpendicular to image plane

Crossing fibers can be located in the plane oriented along the beam propagation direction. Fig. 12 a1-a3 shows plywood structures with different polarity. One layer of fibers is fixed parallel to the image plane, and the other layer is tilted out to the image plane. SHG intensity is high for the crossed layers of parallel fibers (a1) and drops significantly for antiparallel fibers of the layers (a2, a3). SHG intensity increases with the layer tilt and then decreases at large angles for collagen, and decreases with tilt for TPPS_4_ aggregates. The corresponding R ratios (Fig. 12 d and e) have similar initial values to parallel configuration (*σ*=0) without tilt (*α*=0) for collagen and TPPS_4_ fibers (Fig. 10 d and e, respectively). The R ratio increases with the axial tilt and then decreases at large angles for collagen, and slightly decreases and then slightly increases for TPPS_4_ fibers. The C ratio behaviors (Fig. 12, f and g) show typical sin *α* curves. Changing the handedness of individual fibers in the layers results in flipping the sign of C ratio for both structures (Fig. 12 f and g). The dependencies of *I*, R and C on alpha are similar to the axially two crossing fibers in Fig 9.

**Figure 12:**
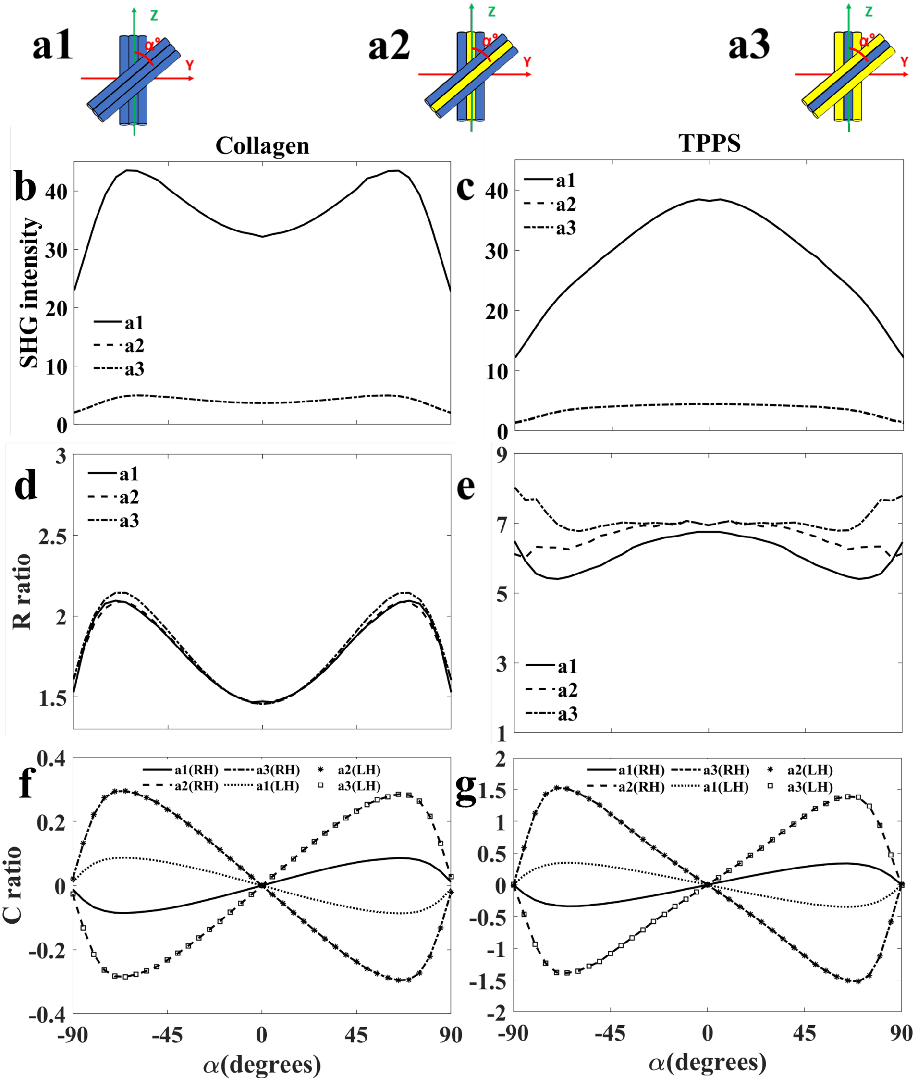
Axially crossing plywood structures with parallel (a1) and antiparallel (a2, a3) crossing layers. One layer is parallel to the image plane, and the other layer is tilted axially by an angle *α*. SHG intensity, *I*, (b, c), R ratio (d, e), and C ratio (f, g) variations with tilt angle *α* for collagen and TPPS_4_ aggregates. The figure legend indicates the corresponding fiber configuration in (a). The changes in C ratio are observed for left handed (LH) and right handed (RH) structures in panels (f, g).

### 3.4 Arbitrary oriented 3D crossing fibers, cone geometry

#### 3.4.1 Cone of crossing fibers

##### 3.4.1.1 Fiber cone with effective axis in the image plane (crossing angle dependence)

For three and more crossing fibers at different angles in 3D, a cone geometry can be applied. The effective cone axis of the fibers can be parallel to the image plane (Fig. 13, a) and the cone half angle between the fibers and the cone axis can increase from 0 to 90°. The effect of cone angle dependence on *I*, R and C obtained from digital PIPO microscope is shown in Fig. 13. The intensity decreases by increasing the cone angle for both collagen and TPPS_4_ fibers (Fig. 13, b and c, respectively). Fig. 13, d and e show that R ratio increases for collagen, and decreases for TPPS_4_ fibers. C ratio decreases from 0 to 45°and then increases with increasing *σ* for both samples (Fig. 13, f, g). Switching the handedness of the fibers results in changing the sign of C ratio in both aggregate types.

**Figure 13:**
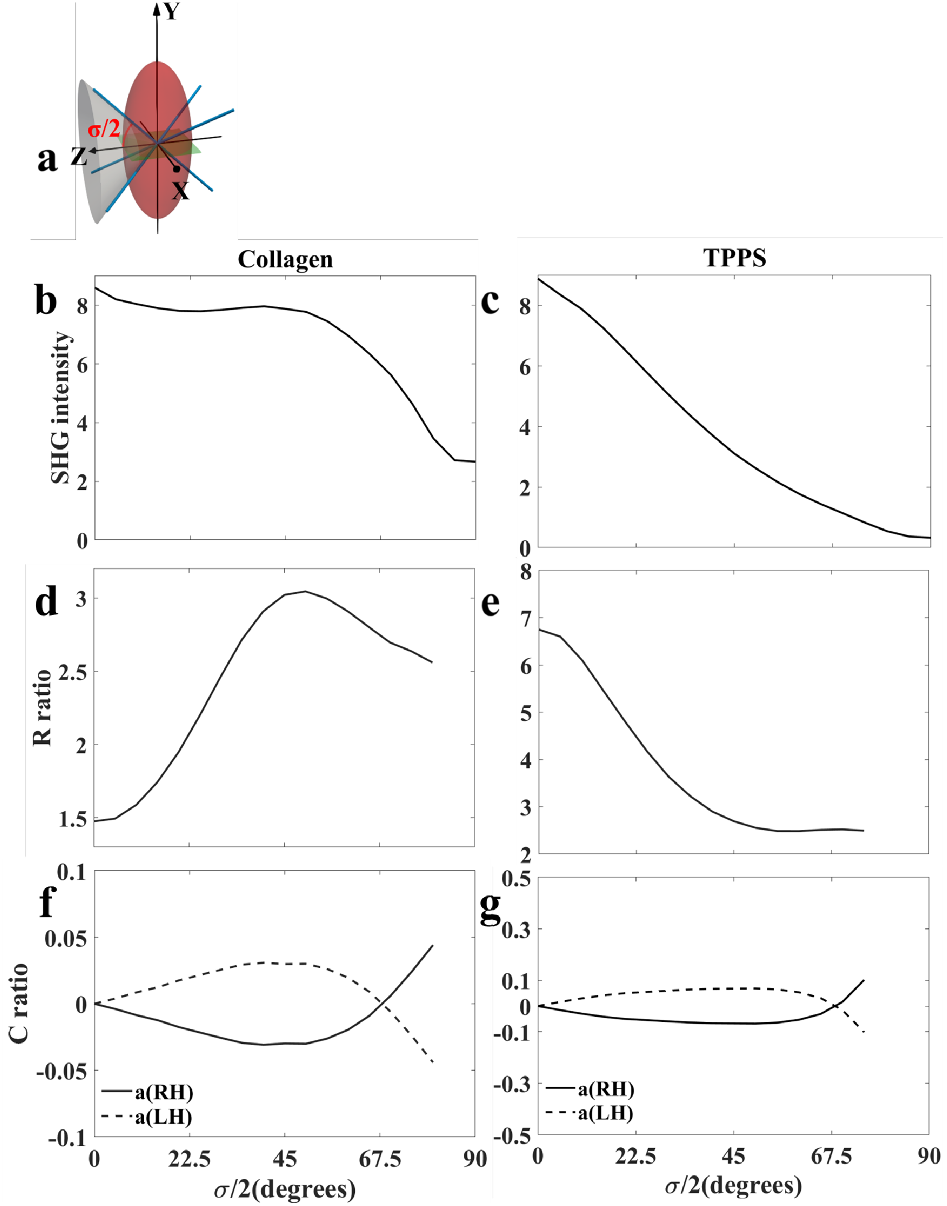
SHG dependence on the half angle *σ/*2 of the cone comprised of three fibers, with effective cone axis in the image plane. Schematic picture of the cone structure with cone axis in the image plane (a). *I* (b, d), R ratio (d, e), and C ratio (f, g) changes with cone angle *σ/*2 for collagen (b, d, f) and TPPS_4_ fibers (c, e, g). Figure legend indicates right handed (RH) and left handed (LH) fibers in panels f and g.

##### 3.4.1.2 Fiber cone with effective axis tilting out of the image plane

The cone axis can be tilted out of the image plane. The *α* dependence of *I*, R, and C ratio for tilting the cone axis is presented in Fig. 14. The cone structure is composed of two fibers with crossing angle of 45°in the image plane (-22.5°and +22.5°respect to Z axis), and both are tilted - 22.5°out of the image plane. The third fiber is initially parallel to the Z axis and then tilted +22.5°out of the image plane. The normalized SHG intensity dependence on *α* is plotted in Fig. 14, b and c for collagen and TPPS_4_ aggregates, respectively. The *I* dependence on *α* has maxima at -45°and +45°for collagen and increases monotonously with *α* for TPPS_4_ aggregates. The R ratio changes are plotted in Fig. 14, d and e, where R peaks at -45°and +45°are observed for collagen, and gradual R increase with *α* for TPPS_4_ fibers. The C ratio changes are presented in Fig. 14, f and g. The sign of C ratio changes by changing the handedness of individual fibers.

**Figure 14:**
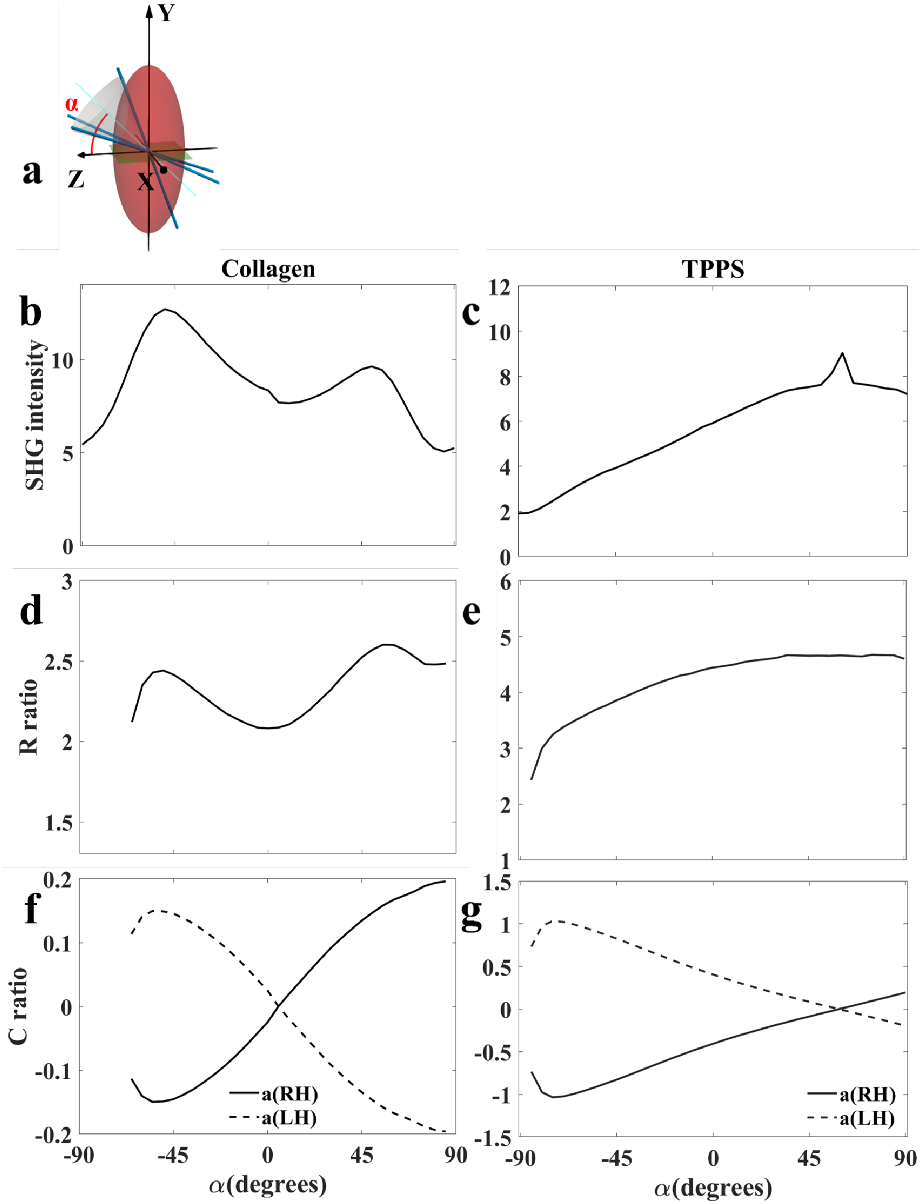
Tilting a 3D cone structure out of the image plane. Schematic picture of a tilted cone structure by an angle *α* from the image plane to the cone axis (a). Dependence of normalized SHG intensity (b, c) R (d, e) and C (f, g) ratio on the tilt angle *α* of the cone axis for collagen (b, d, f) and TPPS_4_ (c, e, g). The left hand (LH) and right hand (RH) fiber structure is indicated in the figure legends (f, g).

#### 3.4.2 Isotropically distributed crossing fibers in 3D

Isotropic structure can be constructed from fibers oriented with equal probability in all 3 dimensions. The SHG intensity is very low in isotropic structures and appears due to Hyper-Rayleigh scattering (35). The PIPO digital microscope calculates *I* = 0.75*I*_0_ for isotropic arrangement of collagen and *I* = 0.4*I*_0_ for TPPS_4_ fibers. The PIPO digital microscope results cannot be analyzed by *C*_6_ symmetry model and therefore R and C ratios are not retrieved.

## 4 Discussion

The impacts of different fiber configurations in the focal volume on *I*, R and C have been investigated numerically with PIPO SHG digital microscope and summarized in Table 1. The fibers are assumed to have C6 symmetry with real valued achiral and complex valued chiral susceptibility components (15, 25). The effects differ for fibers with molecular achiral susceptibility of the fibers below and above the value of 3, representing collagen and TPPS_4_ fibers, respectively. The following effects are revealed:

**Table 1:** Table of the major effects in the focal volume of the PIPO microscope.

|  | Collagen |  |  | TPPS<sub>4</sub> |  |  | Ref. |
| --- | --- | --- | --- | --- | --- | --- | --- |
|  | *I* | *R* | *C* | *I* | *R* | *C* |  |
| **Displacing one fiber laterally** | follows PSF | change with handedness | no change | follows PSF | change with handedness | no change | - |
| **Polarity** | coherent sum | changes for antiparallel | changes for antiparallel | coherent sum | changes for antiparallel | changes for antiparallel | (30–33) |
| **Tilting out of the image plane** | increases | increases | $\propto \sin(\alpha)$ | decreases | decreases | $\propto \sin(\alpha)$ | (19) |
| **Crossing fibers** | decreases | increases | if $\alpha = 0$, $C = 0$ | decreases | decreases | if $\alpha = 0$, $C = 0$ | (2, 19) |
| **Fiber handedness** | no change | no change | sign change | no change | no change | sign change | (28) |
| **Macrochirality** | no change | no change | sign change | no change | no change | sign change | (26) |

1. A novel effect is observed with an individual chiral fiber in a voxel, where R ratio changes when moving the fiber laterally from one to another side in the focal volume (Fig 2). This effect appears only for fibers with a complex valued chiral susceptibility tensor component.
2. The same polarity fibers coherently enhance SHG intensity, and antiparallel fibers strongly diminish the signal (Fig. 4 and 5). The effect on R ratio is less pronounced for different amount of fiber polarity in a structure (Fig. 6). The change of C ratio with tilt depends on the amount and configuration of different polarity fibers, and might have pronounced enhancement of C ratio with tilt for close to equal number of antiparallel fibers (Fig. 6).
3. The tilt of fibers out of image plane increases SHG intensity for collagen and decreases for TPPS_4_ fibers. The R ratio increases with tilt for collagen and decreases for TPPS_4_ (see also eq (2)). The C ratio shows a sinusoidal behavior with tilt, and changes sign for opposite chirality of the fiber. Tilting two parallel fibers out of the image plane provides the same C ratio dependence as for one fiber, but antiparallel fibers cancel each other’s effect upon tilt (Fig. 5). Tilting a bundle of antiparallel fibers may result in high deviation from C ratio dependence described by eq (3) (Fig. 6).
4. The crossing fibers affect SHG intensity, R and C ratios. The SHG intensity decreases with the crossing angle increase. The analytical calculations show that two fibers with similar R and C ratios crossing in the image plane are equivalent to one fiber aligned with an effective orientation and a modified R ratio. The R ratio increases for collagen with increasing the crossing angle, and decreases for TPPS_4_ aggregates. In general, several fibers can cross in the focal volume in 3D arrangement, making approximate cone with effective orientation axis. The R ratio is modified at different crossing angles in the same way as for two crossing fibers. The previous studies suggest simple addition of SHG intensities for crossing fibers (2, 36). The agreement between analytical and numerical calculations in this work shows that addition of SHG electric field amplitudes is sufficient for adjacent crossing fibers, but phase relations have to be included for larger fiber separations.
5. Changing handedness of the fibers does not change SHG intensity and R ratio, but changes the sign of C ratio (Figs. 3, 5, 8, 9, 11, 12, 13, 14). The sign and magnitude of C ratio is influenced by tilt angle and polarity of the fibers.
6. The macrochirality effect can occur in plywood geometry at large crossing angles. SHG intensity and R ratio is not affected, but C ratio has a small effect differing for clockwise and counterclockwise crossed fiber layers (Fig. 10).

A summary of the major SHG polarimetric effects for different fiber arrangements in the focal volume is outlined in Table 1. The geometry of a real biological structure in the focal volume can experience a mixture of the effects. Complex configurations such as tilted and crossed antiparlallel plywood structures and antiparallel bundles are modeled in this work to help in interpreting the PIPO microscopy results.

## 5 Conclusion

In this study, SHG responses of chiral fibers with different configurations in the focal volume are modeled with a PIPO digital microscope. There are several fiber configurations that significantly influence the results of polarimetric nonlinear microscopy: 1. An individual chiral fiber has different R ratio when moved from one side to another side in a voxel. 2. The parallel and antiparallel fibers have large differences in SHG intensity due to coherent summation of SHG intensities generated from different fibers, while R and C ratio is also influenced by the polarity of neighboring fibers. 3. The fiber tilt out of image plane affects SHG intensity, R and C ratio. 4. Crossing fibers also affect SHG intensity, R and C ratios. 5. Handedness of the fibers mostly affect the C ratio, and 6. Macrochirality of fibers in a voxel can affect slightly C ratio at large crossing angles. A combination of different effects outlined above can be observed in the biological structures.

This study contributes to the understanding and interpretation of the imaging results of biological tissue obtained by polarimetric SHG microscopy. The modeling results confirm that polarimetric microscopy is a powerful imaging technique that enables ultrastructural determination of various fibrillar configurations within the voxels, and helps to reconstruct the 3D fiber organization in biological tissues.

## Data availability

The datasets used during the current study are available from the corresponding author on reasonable request.

## References

1. Mansfield, J. C., V. Mandalia, A. Toms, C. P. Winlove, and S. Brasselet, 2019. Collagen reorganization in cartilage under strain probed by polarization sensitive second harmonic generation microscopy. Journal of the Royal Society Interface 16:20180611.

2. Alizadeh, M., D. Merino, G. Lombardo, M. Lombardo, R. Mencucci, M. Ghotbi, and P. Loza-Alvarez, 2019. Identifying crossing collagen fibers in human corneal tissues using pSHG images. Biomedical Optics Express 10:3875–3888.

3. Golaraei, A., K. Mirsanaye, Y. Ro, S. Krouglov, M. K. Akens, B. C. Wilson, and V. Barzda, 2019. Collagen chirality and three-dimensional orientation studied with polarimetric second-harmonic generation microscopy. Journal of biophotonics 12:e201800241.

4. Chen, X., O. Nadiarynkh, S. Plotnikov, and P. J. Campagnola, 2012. Second harmonic generation microscopy for quantitative analysis of collagen fibrillar structure. Nature protocols 7:654–669.

5. Zipfel, W. R., R. M. Williams, R. Christie, A. Y. Nikitin, B. T. Hyman, and W. W. Webb, 2003. Live tissue intrinsic emission microscopy using multiphoton-excited native fluorescence and second harmonic generation. Proceedings of the National Academy of Sciences 100:7075–7080.

6. Plotnikov, S. V., A. C. Millard, P. J. Campagnola, and W. A. Mohler, 2006. Characterization of the myosin-based source for second-harmonic generation from muscle sarcomeres. Biophysical journal 90:693–703.

7. Tiaho, F., G. Recher, and D. Rouède, 2007. Estimation of helical angles of myosin and collagen by second harmonic generation imaging microscopy. Optics express 15:12286–12295.

8. Alizadeh, M., M. Ghotbi, P. Loza-Alvarez, and D. Merino, 2019. Comparison of Different Polarization Sensitive Second Harmonic Generation Imaging Techniques. Methods and Protocols 2:49.

9. Psilodimitrakopoulos, S., I. Amat-Roldan, P. Loza-Alvarez, and D. Artigas, 2012. Effect of molecular organization on the image histograms of polarization SHG microscopy. Biomedical optics express 3:2681–2693.

10. Psilodimitrakopoulos, S., I. Amat-Roldan, P. Loza-Alvarez, and D. Artigas, 2010. Estimating the helical pitch angle of amylopectin in starch using polarization second harmonic generation microscopy. Journal of Optics 12:084007.

11. Chu, S.-W., S.-Y. Chen, G.-W. Chern, T.-H. Tsai, Y.-C. Chen, B.-L. Lin, and C.-K. Sun, 2004. Studies of χ (2)/χ (3) tensors in submicron-scaled bio-tissues by polarization harmonics optical microscopy. Biophysical journal 86:3914–3922.

12. Psilodimitrakopoulos, S., S. I. Santos, I. Amat-Roldan, T. K. Nair, D. Artigas-García, and P. Loza-Alvarez, 2009. In vivo, pixel-resolution mapping of thick filaments’ orientation in nonfibrilar muscle using polarization-sensitive second harmonic generation microscopy. Journal of Biomedical Optics 14:014001.

13. Stoller, P., K. M. Reiser, P. M. Celliers, and A. M. Rubenchik, 2002. Polarization-modulated second harmonic generation in collagen. Biophysical journal 82:3330–3342.

14. Amat-Roldan, I., S. Psilodimitrakopoulos, P. Loza-Alvarez, and D. Artigas, 2010. Fast image analysis in polarization SHG microscopy. Optics express 18:17209–17219.

15. Golaraei, A., L. Kontenis, K. Mirsanaye, S. Krouglov, M. K. Akens, B. C. Wilson, and V. Barzda, 2019. Complex susceptibilities and chiroptical effects of collagen measured with polarimetric second-harmonic generation microscopy. Scientific reports 9:1–12.

16. Samim, M., S. Krouglov, and V. Barzda, 2015. Double Stokes Mueller polarimetry of second-harmonic generation in ordered molecular structures. JOSA B 32:451–461.

17. Psilodimitrakopoulos, S., P. Loza-Alvarez, and D. Artigas, 2014. Fast monitoring of in-vivo conformational changes in myosin using single scan polarization-SHG microscopy. Biomedical optics express 5:4362–4373.

18. Samim, M., S. Krouglov, and V. Barzda, 2016. Nonlinear Stokes-Mueller polarimetry. Physical Review A 93:013847.

19. Tuer, A. E., M. K. Akens, S. Krouglov, D. Sandkuijl, B. C. Wilson, C. M. Whyne, and V. Barzda, 2012. Hierarchical model of fibrillar collagen organization for interpreting the second-order susceptibility tensors in biological tissue. Biophysical journal 103:2093–2105.

20. Golaraei, A., R. Cisek, S. Krouglov, R. Navab, C. Niu, S. Sakashita, K. Yasufuku, M.-S. Tsao, B. C. Wilson, and V. Barzda, 2014. Characterization of collagen in non-small cell lung carcinoma with second harmonic polarization microscopy. Biomedical optics express 5:3562–3567.

21. Tokarz, D., R. Cisek, A. Golaraei, S. Krouglov, R. Navab, C. Niu, S. Sakashita, K. Yasufuku, M.-S. Tsao, S. L. Asa, et al., 2015. Tumor tissue characterization using polarization-sensitive second harmonic generation microscopy. In Bio-photonics South America. International Society for Optics and Photonics, volume 9531, 95310C.

22. Tokarz, D., R. Cisek, A. Joseph, S. L. Asa, B. C. Wilson, and V. Barzda, 2020. Characterization of pathological thyroid tissue using polarization-sensitive second harmonic generation microscopy. Laboratory Investigation 100:1280–1287.

23. Tuer, A. E., S. Krouglov, N. Prent, R. Cisek, D. Sandkuijl, K. Yasufuku, B. C. Wilson, and V. Barzda, 2011. Nonlinear optical properties of type I collagen fibers studied by polarization dependent second harmonic generation microscopy. The Journal of Physical Chemistry B 115:12759–12769.

24. Sandkuijl, D., A. E. Tuer, D. Tokarz, J. Sipe, and V. Barzda, 2013. Numerical second-and third-harmonic generation microscopy. JOSA B 30:382–395.

25. Schmeltz, M., C. Teulon, M. Pinsard, U. Hansen, M. Alnawaiseh, D. Ghoubay, V. Borderie, G. Mosser, C. Aimé, F. Légaré, et al., 2020. Circular dichroism second-harmonic generation microscopy probes the polarity distribution of collagen fibrils. Optica 7:1469–1476.

26. Abramavicius, D., S. Krouglov, and V. Barzda, 2021. Second harmonic generation theory for a helical macromolecule with high sensitivity to structural disorder. Physical Chemistry Chemical Physics 23:20201–20217.

27. Simpson, G. J., and K. L. Rowlen, 1999. An SHG magic angle: dependence of second harmonic generation orientation measurements on the width of the orientation distribution. Journal of the American Chemical Society 121:2635–2636.

28. Pleckaitis, M., F. Habach, L. Kontenis, G. Steinbach, G. Jarockyte, A. Kalnaityte, I. Domonkos, P. Akhtar, M. Alizadeh, S. Bagdonas, et al., 2022. Structure and principles of self-assembly of giant “sea urchin” type sulfonatophenyl porphine aggregates. Nano Research 1–11.

29. Lombardo, M., D. Merino, P. Loza-Alvarez, and G. Lombardo, 2015. Translational label-free nonlinear imaging biomarkers to classify the human corneal microstructure. Biomedical Optics Express 6:2803–2818.

30. Pfeffer, C. P., B. R. Olsen, F. Ganikhanov, and F. Légaré, 2008. Multimodal nonlinear optical imaging of collagen arrays. Journal of structural biology 164:140–145.

31. Harnagea, C., M. Vallières, C. P. Pfeffer, D. Wu, B. R. Olsen, A. Pignolet, F. Légaré, and A. Gruverman, 2010. Two-dimensional nanoscale structural and functional imaging in individual collagen type I fibrils. Biophysical journal 98:3070–3077.

32. Bancelin, S., C.-A. Couture, K. Légaré, M. Pinsard, M. Rivard, C. Brown, and F. Légaré, 2016. Fast interferometric second harmonic generation microscopy. Biomedical optics express 7:399–408.

33. Stoller, P., P. M. Celliers, K. M. Reiser, and A. M. Rubenchik, 2003. Quantitative second-harmonic generation microscopy in collagen. Applied optics 42:5209–5219.

34. Burke, M., A. Golaraei, A. Atkins, M. Akens, V. Barzda, and C. Whyne, 2017. Collagen fibril organization within rat vertebral bone modified with metastatic involvement. Journal of structural biology 199:153–164.

35. Hollis, D. B., 1988. Review of hyper-Rayleigh and second-harmonic scattering in minerals and other inorganic solids. American Mineralogist 73:701–706.

36. Latour, G., I. Gusachenko, L. Kowalczuk, I. Lamarre, and M.-C. Schanne-Klein, 2012. In vivo structural imaging of the cornea by polarization-resolved second harmonic microscopy. Biomedical optics express 3:1–15.

